# Gene-centered identification of cis-regulatory islands reveals regulatory landscapes complementary to motif-centric approaches

**DOI:** 10.64898/2026.01.05.697455

**Authors:** Yoshihiro Ohmori

## Abstract

Cis-regulatory elements constitute a fundamental layer of gene regulation, yet their computational identification has largely relied on transcription factor (TF)–centric frameworks that assume genome-wide background normalization and explicit TF binding models. While effective at the genome scale, such assumptions are less appropriate for gene-centered analyses, where local sequence composition rather than global averages defines the relevant regulatory context. Here, we introduce a TF-independent framework for the gene-centered identification of cis-regulatory islands (GCIC), which detects regulatory structure based on the local enrichment and diversity of short cis-regulatory sequence words derived from curated plant regulatory elements. Cis-regulatory islands are identified through the spatial overlap of independently enriched motif families, without relying on TF identity, binding affinity, or genome-wide normalization. Application of the GCIC framework to the *DROOPING LEAF* (*DL*) locus in rice identifies discrete cis-regulatory islands, including one that coincides with a previously characterized intronic regulatory region, and reveals spatial patterns distinct from those detected by PWM-based motif scanning and motif clustering approaches. Genome-wide analyses further show that cis-regulatory islands are broadly distributed across genes but exhibit heterogeneous motif-family usage: regulatory vocabulary diversity expands at the gene level, whereas individual islands preferentially reuse a limited set of motif-family combinations. These results indicate that cis-regulatory organization is best described as a gene-centered property of sequence vocabulary usage, in which regulatory diversity arises through gene-specific deployment and constrained reuse of motif-family combinations rather than unrestricted combinatorial complexity. The GCIC framework thus provides a complementary representation of regulatory landscapes tailored to gene-centered analyses, capturing regulatory features that are not readily detected by motif-centric approaches optimized for genome-wide inference.

## Introduction

Cis-regulatory elements constitute a fundamental layer of gene expression control and underlie the spatial, temporal, and quantitative regulation of transcription (Levine and Tjian, 2003; Wittkopp and Kalay, 2011). Over the past decades, computational identification of cis-regulatory regions has been dominated by transcription factor (TF)–centric frameworks, in which regulatory sequences are inferred by scanning genomes for matches to position weight matrices (PWMs) representing TF binding preferences (Stormo, 2000; Wasserman and Sandelin, 2004). Detected motif hits are then aggregated or clustered to define cis-regulatory modules (CRMs), an approach that has proven highly effective in genome-wide analyses where appropriate background models suppress spurious enrichment (Frith et al., 2003; Bailey et al., 2009; Grant et al., 2011).

Despite their success, TF-centric methods rest on implicit assumptions about regulatory organization. Most notably, they assume that cis-regulatory structure is primarily defined by TF identity and that genome-wide background normalization provides an appropriate null model for motif occurrence, implicitly treating genome-wide sequence composition as a regulatory-neutral baseline (Spitz and Furlong, 2012). These assumptions become less appropriate when the analytical focus shifts from genome-averaged patterns to gene-centered questions, such as characterizing regulatory architecture specific to individual loci. Under such gene-centered conditions, motif density–based approaches can become dominated by sequence features that are typically suppressed at the genome scale, including low-complexity or repetitive sequence composition, thereby obscuring gene-specific regulatory organization rather than revealing it (Tautz et al., 1986; Hardison and Taylor, 2012).

More broadly, existing cis-regulatory sequence analysis frameworks differ substantially in their primary observables, analytical units, and underlying assumptions regarding how regulatory information is encoded and organized in genomic sequences. A conceptual comparison of representative experimental and computational frameworks is provided in Supplementary Table S1.

An alternative perspective is that cis-regulatory organization may be encoded in the local distribution and combination of short regulatory sequence words, independent of explicit TF identity. From this viewpoint, transcription factors act as interpreters of pre-existing regulatory sequence landscapes rather than as the sole architects of regulatory structure (Levine and Tjian, 2003). Accordingly, regulatory organization cannot be fully captured by motif rarity or TF identity alone, but must be defined relative to the local sequence environment of individual genes. Gene-centered analyses that focus on local enrichment and diversity of regulatory sequence vocabulary may therefore reveal regulatory structure that is complementary to, but not reducible to, PWM-based motif clustering.

Plant cis-regulatory element databases such as PLACE provide curated collections of experimentally characterized regulatory sequences, many of which consist of short and degenerate motifs typically on the order of 4–8 bp (Higo et al., 1999). While biologically meaningful, these motifs do not necessarily correspond to high-information-content transcription factor binding sites and are therefore not optimally exploited by standard PWM-based scanning and clustering frameworks, particularly under gene-centered analytical conditions.

Here, we introduce a TF-independent, gene-centered framework for the gene-centered identification of cis-regulatory islands (GCIC), based on the local enrichment and diversity of short cis-regulatory sequence words derived from curated plant regulatory elements. By defining enrichment relative to predefined gene-centered genomic regions and integrating signals across motif families, the GCIC framework identifies cis-regulatory islands without explicit modeling of transcription factor binding. Direct comparison with PWM-based motif scanning (FIMO) and motif clustering (Cluster-Buster) under identical gene-centered conditions demonstrates that GCIC captures discrete regulatory landscapes consistent with known transcriptional regulation, while revealing regulatory organization that remains obscured under motif-centric approaches. An overview of the conceptual framework and the definition of cis-regulatory islands under GCIC are illustrated in Figure 1.

**Figure 1.**
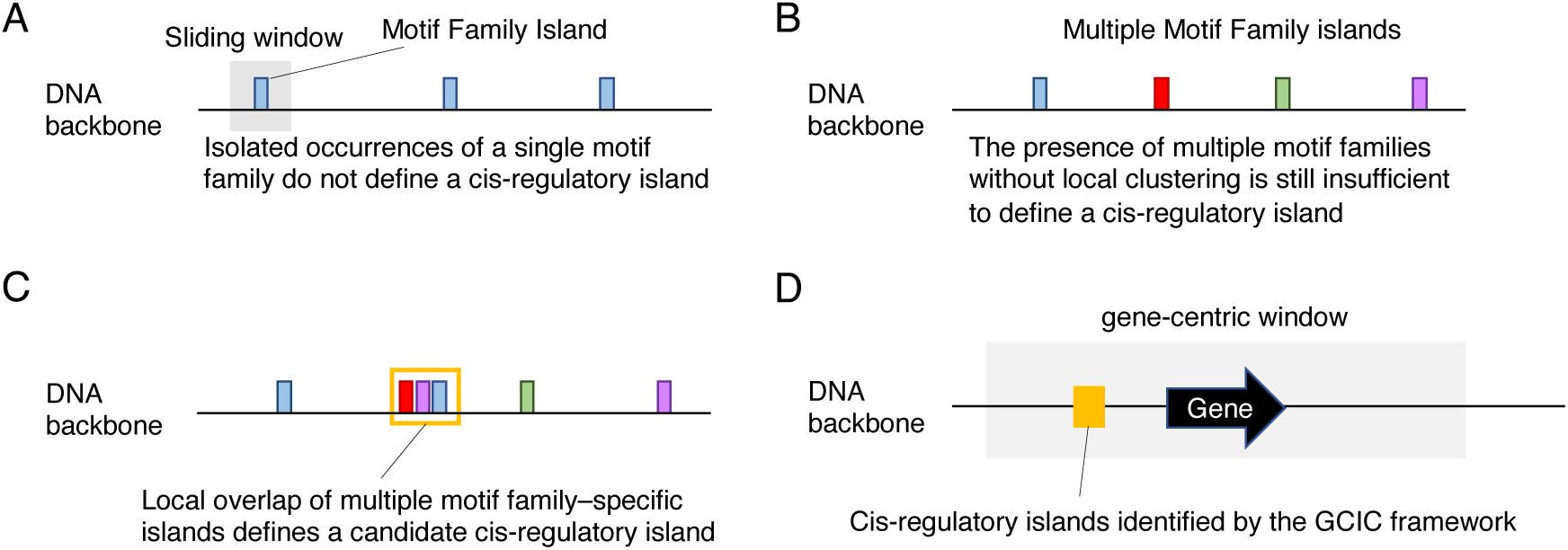
| Conceptual framework of gene-centered identification of cis-regulatory islands (GCIC) (A) Detection of motif family–specific enriched regions. Occurrences of short cis-regulatory sequence words are grouped into motif families and quantified along the genomic sequence using a sliding-window approach. Family-wise enrichment is evaluated locally using a slidingwindow–based Z-score framework. (B) Enrichment of individual motif families alone is insufficient to define regulatory structure when enriched regions are spatially dispersed and do not overlap within a genomic interval. (C) Local spatial overlap among independently enriched motif families defines a candidate cisregulatory island. (D) A predefined gene-centered genomic window is specified for each gene. Candidate regions identified within this window by the GCIC framework are operationally defined as cisregulatory islands.

## Methods

### Definition of cis-regulatory sequence vocabulary derived from PLACE

Cis-regulatory sequence vocabularies were derived from the PLACE database (version 30.0), which comprises 469 experimentally reported cis-acting regulatory DNA motif entries curated from the literature. PLACE was originally developed to cover vascular plants and was subsequently expanded to include motif variants reported across multiple plant species, reflecting the accumulation of regulatory motif evidence beyond individual genes or taxa.

In addition to canonical motif definitions, PLACE includes documented sequence variations identified in different genes and plant species, along with associated literature references and nucleotide sequence accession information. As such, PLACE represents a cross-species cis-regulatory vocabulary rather than a species-specific motif collection, making it suitable for comparative and gene-centered analyses across plant genomes.

PLACE motifs include both exact and degenerate consensus sequences represented using IUPAC nucleotide codes and were used as provided without trimming, k-mer enumeration, or conversion into position weight matrices. Each motif sequence was converted into an IUPAC-expanded regular expression and scanned directly against genomic sequences. At no stage of the analysis were transcription factor identities, binding affinities, or PWM-based models assumed or inferred; detected motif occurrences were treated strictly as sequence features rather than as predictions of transcription factor binding events.

For downstream analyses, individual motif occurrences were grouped into motif families defined in this study. Motif family assignments were manually curated based on sequence similarity and reported regulatory usage, and the corresponding mappings are provided in Supplementary Data 1 (family_map.tsv). Enrichment analyses were performed at the motif family level rather than for individual motifs.

PLACE motifs vary substantially in length and sequence complexity, ranging from short elements of approximately 4–8 bp to longer consensus sequences containing degenerate bases or low-complexity features. In this respect, PLACE motifs constitute a cis-regulatory vocabulary that is fundamentally distinct from the longer, transcription factor–specific motifs typically assumed in PWM-based motif-centric analyses. Motif scanning was performed on strand-normalized genomic sequences using PLACE motif definitions without explicit enumeration of reverse-complement sequences. Consequently, detected motif family usage represents a conservative estimate.

### Construction of gene-centered genomic regions

Gene-centered genomic regions were constructed using the *Oryza sativa* IRGSP-1.0 reference genome and gene annotation obtained from Ensembl Plants (release 58; files: Oryza_sativa.IRGSP-1.0.dna_sm.toplevel.fa and Oryza_sativa.IRGSP-1.0.58.gtf).

For each annotated gene, the gene body defined in the GTF file was first treated as a genomic block. Upstream and downstream regions were then extended from this block up to a maximum of 10 kb on each side, subject to predefined genomic window constraints. Regions that were shorter than the minimum required length or that contained a high fraction of undefined nucleotides (N) were excluded based on quality-control criteria.

For downstream sequence-based analyses, all retained gene-centered sequences were strand-normalized to preserve the 5′–3′ transcriptional orientation. Sequences derived from genes annotated on the negative strand were reverse-complemented prior to analysis. Genomic coordinates and basic properties of all gene-centered regions used in this study are provided in Supplementary_Data 2 (gene_regions.tsv).

All subsequent genome-wide analyses were restricted to these predefined gene-centered regions to ensure that comparisons among methods were performed on identical genomic intervals and were not influenced by genome-wide background normalization.

### Detection of cis-regulatory islands using the gene-centered identification framework

Cis-regulatory islands were detected using the Gene-centered identification of cis-regulatory islands (GCIC) framework, which evaluates local enrichment of PLACE-derived motif families within predefined gene-centered genomic regions.

Each gene-centered region was scanned using sliding windows of 200 bp with a step size of 50 bp. Within each window, occurrences of PLACE motifs were counted and aggregated according to motif family.

For each motif family, window-level counts were normalized by Z-score transformation across all windows within the same gene-centered region. This normalization was performed independently for each gene, thereby defining enrichment relative to the local sequence context of that gene rather than to a genome-wide background distribution. Windows with Z-scores ≥ 2.5 were considered to show significant local enrichment for the corresponding motif family.

Adjacent or overlapping enriched windows belonging to the same motif family were merged to define family-specific cis-regulatory islands. Regions in which cis-regulatory islands from two or more distinct motif families spatially overlapped were then retained as cis-regulatory islands identified under the GCIC framework. Only regions supported by such multi-family overlap were retained; outside these regions, combinatorial enrichment was not evaluated by definition.

Under the GCIC framework, cis-regulatory organization is identified as a property of local sequence composition conditioned on individual genes. This approach does not assume transcription factor binding, motif clustering density, predefined regulatory module length, or genome-wide background normalization, and instead characterizes regulatory structure through the coordinated enrichment of diverse regulatory sequence vocabularies within gene-centered contexts.

### Locus-specific analysis of the *DL* region

For illustrative purposes, the *DROOPING LEAF* (*DL*) locus was analyzed separately using a contiguous genomic window spanning approximately 35 kb. In this locus-specific analysis, cis-regulatory island detection was performed using the same GCIC framework, motif family definitions, sliding window parameters, and Z-score thresholds as described above. However, normalization was applied across sliding windows within the locus-wide region rather than within gene-centered windows.

This locus-specific analysis was used solely for visualization and conceptual illustration of cis-regulatory island organization at a well-characterized regulatory locus and was not included in genome-wide statistical summaries or gene-level enrichment analyses.

### PWM-based motif scanning and clustering

For comparison with motif-centric approaches, PWM-based motif scanning was performed using FIMO (Grant et al., 2011) with default parameters on the *DL* locus, using the same contiguous 35 kb genomic region analyzed under the GCIC framework. Motif hit densities were summarized along this locus-wide region and visualized using both linear and log10-scaled representations.

Motif clustering was performed using Cluster-Buster (Frith et al., 2003) with standard settings to identify cis-regulatory modules (CRMs) characterized by dense clusters of PWM hits within the same 35 kb region. No repeat masking or genome-wide background normalization was applied in these analyses, in order to directly assess the behavior of PWM-based methods under identical locus-scale conditions.

### Extraction of cis-regulatory region sequences

To inspect internal sequence composition, genomic sequences corresponding to cis-regulatory islands identified under the GCIC framework and CRMs were extracted directly from the gene-centered FASTA sequences used as input. Sequence extraction was performed using bedtools getfasta (Quinlan and Hall, 2010), without repeat masking or additional filtering. Extracted sequences were used solely for descriptive examination of internal sequence features and did not influence the definition or boundaries of cis-regulatory islands or CRMs.

### Genome-wide application, quantification, and annotation of GCIC islands

The GCIC framework was applied genome-wide to all annotated genes using the same parameters, motif family definitions, sliding window settings, and gene-centered genomic regions described above. For each gene, the presence or absence of cis-regulatory islands identified under the GCIC framework, the number of such islands, and their genomic coordinates were recorded.

Genome-wide GCIC annotations were subsequently used for quantitative analyses at both the gene and cis-regulatory island levels. For each gene, motif-family usage and regulatory vocabulary diversity were quantified based on all associated GCIC islands. At the cis-regulatory island level, motif-family vocabularies were defined as the unique combinations of motif families present within individual islands, and vocabulary usage frequencies were calculated across the genome.

The genomic context of GCIC islands was determined by intersecting island coordinates with gene annotations from Ensembl Plants. GCIC islands were classified according to overlap with promoter, exonic, intronic, or intergenic regions. Promoter regions were defined as ±1 kb windows centered on annotated transcription start sites, with strand orientation taken into account.

Gene-level summaries of GCIC island occurrence and motif-family usage are provided in Supplementary Table S2, motif-family vocabulary frequencies across GCIC islands are provided in Supplementary Table S3, and genome-wide GCIC island annotations with genomic context classifications are provided in Supplementary Data 3.

### Integration of GCICs with conserved and functional cis-regulatory datasets

To position GCICs within the established cis-regulatory landscape of rice, GCIC coordinates originally defined on the IRGSP-1.0 reference genome were integrated with conserved and functional cis-regulatory datasets defined on the Azucena genome (AzucenaRS1). The IRGSP-1.0 reference genome sequence was obtained from Ensembl Plants (release 58; file: Oryza_sativa.IRGSP-1.0.dna_sm.toplevel.fa). The Azucena genome sequence was obtained from Ensembl Plants (release 62; file: Oryza_sativa_azucena.AzucenaRS1.dna_sm.toplevel.fa). Because these datasets were generated using different reference genomes, coordinate conversion was required prior to integration.

Whole-genome alignment between the IRGSP-1.0 and Azucena genomes was performed using minimap2 (v2.18; Li, 2018) with parameters optimized for intra-species genome comparison. Based on this alignment, GCIC coordinates were projected from IRGSP-1.0 to the Azucena genome. Only primary, one-to-one syntenic alignment blocks were retained, and GCICs were conservatively lifted over by requiring that each GCIC be fully contained within a single syntenic block. GCICs that did not satisfy this criterion were excluded from downstream analyses. This alignment-based coordinate projection was performed independently of gene annotations, transcription start sites, or transcript models, consistent with the sequence-based and gene-independent design of the GCIC framework.

Genomic coordinates of conserved noncoding sequences (CNSs), transcriptionally active regulatory elements (dREG peaks), and small RNA‒associated genomic features were obtained from the Supplementary Data of Goliasse et al. (2025). These datasets were used as provided, without redefinition, and were analyzed solely as genomic coordinate features without restriction to gene-centered regions.

GCICs projected onto the Azucena genome were used as the reference set for overlap analyses. Overlap between a GCIC and an external regulatory feature was defined as the presence of at least one base pair of shared genomic coordinates. GCICs were classified according to whether they overlapped none, one, or multiple external datasets. In addition, pairwise overlaps among CNSs, dREG peaks, and small RNA‒associated genomic features were quantified to establish a reference framework for interpreting GCIC overlap frequencies.

## Results

### Identification of cis-regulatory islands at a known regulatory locus using the GCIC framework

To illustrate the behavior of the Gene-centered identification of cis-regulatory islands (GCIC) framework, we first applied this approach to the *DROOPING LEAF* (*DL*) locus, for which a regulatory region within the second intron has been experimentally characterized (Ohmori et al., 2011). This locus therefore provides a biologically grounded example for examining how cis-regulatory islands emerge under gene-centered sequence conditioning.

Within the predefined gene-centered genomic region, the analysis identified two cis-regulatory islands under the GCIC framework (Figure 2A). One island was located proximal to the annotated transcription start site, whereas the second island coincided spatially with the previously reported regulatory region in the second intron (Figure 2B). The intronic cis-regulatory island overlapped the experimentally validated cis-regulatory region (CNS-E), demonstrating that cis-regulatory islands identified by the GCIC framework—based on local enrichment of PLACE-derived motif families—can coincide with known regulatory regions despite being detected without transcription factor identity or binding models.

**Figure 2.**
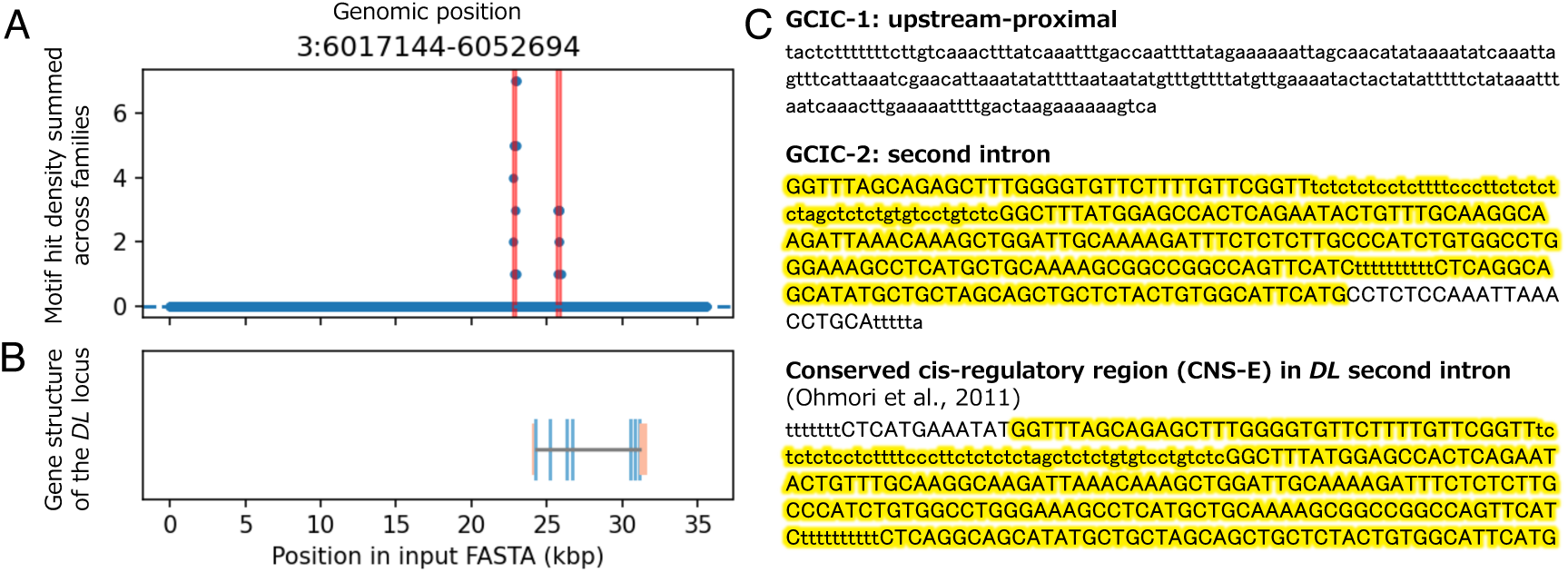
| Identification of cis-regulatory islands at the *DL* locus using the GCIC framework. (**A**) Multi-family motif hit density evaluated along a predefined gene-centered genomic region spanning the *DROOPING LEAF (DL)* locus using a sliding-window approach. Each dot represents a sliding window, and the y-axis indicates the total number of motif hits summed across all motif families within that window. Enrichment is evaluated only within regions that satisfy the criteria for cis-regulatory island detection under the GCIC framework; outside these regions, values are shown as zero for visualization purposes. Red shaded regions indicate cis-regulatory islands identified based on local spatial overlap among independently enriched motif families within the gene-centered region. **(B)** Gene structure of the *DL* locus corresponding to the region shown in (A). Exons are shown as boxes and introns as connecting lines. The positions of cis-regulatory islands relative to gene architecture are indicated to provide gene-centered context. **(C)** Genomic interval–level comparison between cis-regulatory islands identified at the *DL* locus and a previously defined conserved cis-regulatory region (CNS-E) located in the second intron of *DL* (Ohmori et al., 2011). Two cis-regulatory islands were detected within the gene-centered region: an upstream-proximal island located near the annotated transcription start site, and an intronic island. The overlapping genomic interval between the intronic cis-regulatory island and CNS-E is highlighted in yellow on both tracks. Highlighted regions indicate concordance at the level of genomic position rather than base-level sequence identity. Cis-regulatory islands were identified solely based on gene-centered sequence enrichment criteria, without reference to prior functional annotations.

The upstream-proximal cis-regulatory island was not associated with previously reported functional evidence and is therefore presented as a structural detection example rather than as a validated regulatory element. Together, these observations indicate that cis-regulatory islands identified under the GCIC framework emerge as gene-centered regulatory structures reflecting local sequence organization, without presupposing comprehensive functional validation at each locus.

### Gene-centered comparison highlights divergent assumptions among cis-regulatory detection methods

We next compared three cis-regulatory detection strategies—PWM-based motif scanning (FIMO), PWM-based motif clustering (Cluster-Buster), and cis-regulatory island detection using the GCIC framework—using identical gene-centered genomic regions and the same underlying PLACE-derived motif sequences. This gene-centered comparison is summarized in Figure 3. This controlled design removes genome-wide background normalization and isolates how distinct modeling assumptions shape regulatory signal detection under gene-centered constraints.

**Figure 3.**
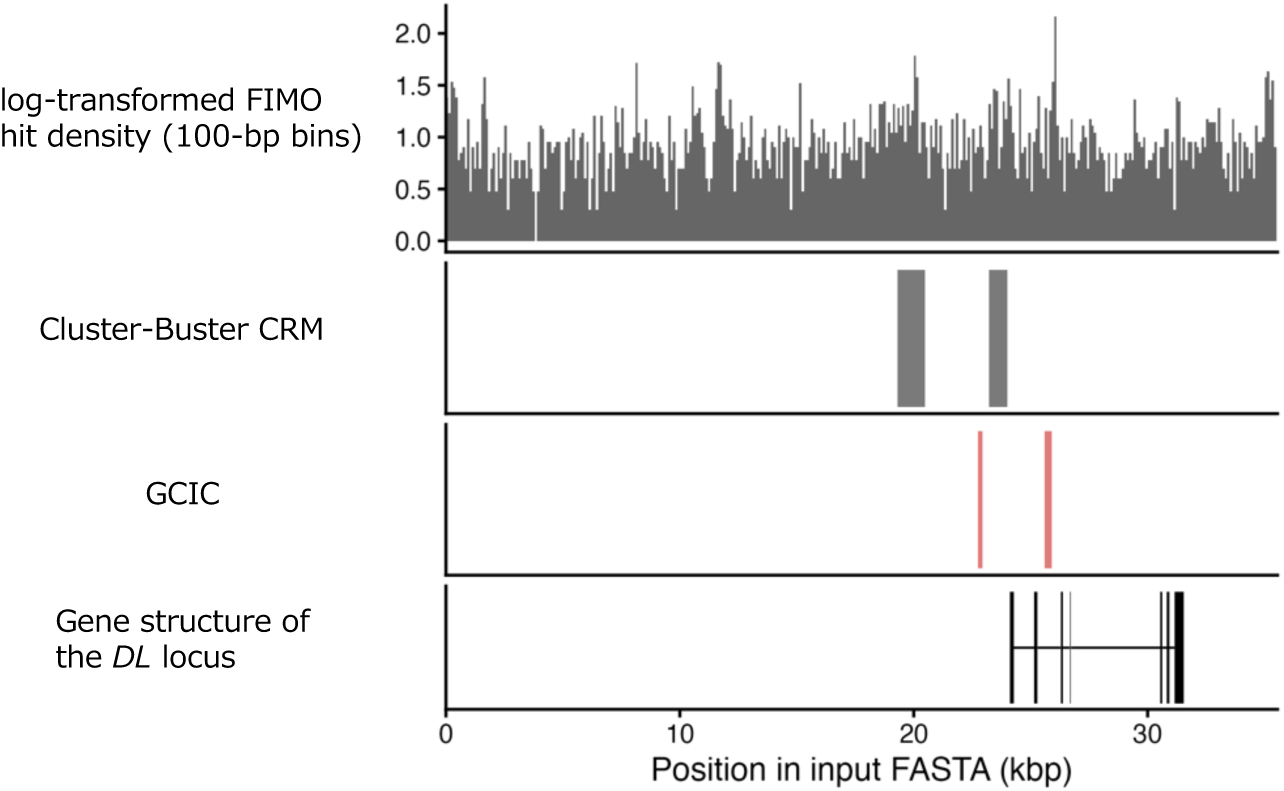
| Gene-centered comparison between motif-centric and cis-regulatory island–based representations. All analyses are shown within the same predefined gene-centered genomic window encompassing the *DROOPING LEAF (DL)* locus, using an identical set of PLACE-derived regulatory motifs. The FIMO track shows the density of PWM-based motif hits summarized in 100-bp bins and visualized as log10(density + 1). Motif occurrences are broadly distributed across the region, producing diffuse signal profiles that largely reflect local sequence composition rather than discrete regulatory units. The Cluster-Buster track displays cis-regulatory modules (CRMs) identified as dense clusters of PWM hits using the same motif set. Detected CRMs span relatively broad genomic intervals and are strongly influenced by local low-complexity and repetitive sequence composition under genecentered conditions without genome-wide background normalization. The GCIC track shows cisregulatory islands identified using the Gene-centered identification of cis-regulatory islands framework. These islands are spatially confined regions defined by the local overlap of independently enriched motif families within the gene-centered window, reflecting regulatory sequence vocabulary diversity rather than motif density or clustering alone. Gene models are shown for reference to indicate the gene-centered coordinate system and to facilitate interpretation of regulatory organization relative to gene architecture.

Consistent with its motif-centric design, FIMO detected numerous motif occurrences broadly distributed across the gene-centered region (Figure 3, FIMO track). When summarized as motif density profiles, these signals exhibited gradual spatial variation that primarily reflected local sequence composition. Log10 transformation compressed the dynamic range but did not resolve discrete regulatory units.

Cluster-Buster identified two prominent cis-regulatory modules (CRMs) characterized by dense motif clustering (Figure 3, Cluster-Buster track). Inspection of the extracted CRM sequences revealed enrichment of short repetitive elements and low-complexity patterns, including GA– and AT-rich repeats (Supplementary Data 4). Such sequences are well known to influence computational motif analyses when background normalization is absent (Tautz et al., 1986). Under the gene-centered conditions applied here, these compositional features dominated the clustering signal, resulting in broad CRM intervals.

In contrast, the GCIC framework identified spatially confined cis-regulatory islands that were distinct from Cluster-Buster CRMs and more localized than the diffuse FIMO signal (Figure 3, GCIC track). These cis-regulatory islands were defined by sustained local enrichment across multiple motif families, reflecting diversity of regulatory sequence vocabulary rather than accumulation of repetitive elements. Notably, cis-regulatory islands were detected even within sequence contexts that include low-complexity segments, indicating that motif family diversity enables discrimination of regulatory structure without explicitly filtering repetitive sequences.

### TF-independent cis-regulatory islands capture gene-centered regulatory organization

Although cis-regulatory islands identified under the GCIC framework are defined without reference to transcription factor identity, their genomic positions were broadly consistent with regions implicated in transcriptional regulation by prior experimental studies, including the intronic regulatory region of *DL* (Ohmori et al., 2011). This concordance supports the view that transcription factor binding operates within an underlying regulatory sequence landscape defined by local sequence composition (Levine and Tjian, 2003; Spitz and Furlong, 2012).

From this perspective, cis-regulatory islands identified using the GCIC framework represent gene-centered regulatory organization rather than predictions of individual TF binding events. The results indicate that regulatory structure can be detected as a property of sequence vocabulary usage conditioned on individual genes, independent of genome-wide statistical assumptions.

Similar positional concordance between GCIC-defined cis-regulatory islands and previously characterized regulatory regions was also observed at additional rice loci. At the *Oryza sativa MADS-box gene 1 (OsMADS1)* locus, a GCIC-defined cis-regulatory island overlaps with an intragenic regulatory region previously shown to be required for proper spatial expression of *OsMADS1*. This regulatory region spans the large first intron and adjacent exonic sequences, which have been experimentally demonstrated to be necessary for flower-preferential expression (Jeon et al., 2008). The GCIC island identified in this study intersects this intragenic interval, indicating concordance at the level of genomic position. A comparable overlap was observed at the *SULPHITE REDUCTASE (SiR)* locus, where a GCIC-defined cis-regulatory island overlaps the distal cis-regulatory module located upstream of the *SiR* coding region. This experimentally characterized module spans approximately − 980 to − 700 bp relative to the translational start site and is required for bundle sheath–specific expression (Hua et al., 2025). The GCIC island overlapped this interval within the gene-centered genomic window used in the present analysis.

These examples are presented as illustrative cases and do not imply comprehensive validation across rice genes. Rather, they indicate that cis-regulatory islands identified by the GCIC framework can coincide with experimentally implicated regulatory regions beyond a single locus, supporting the interpretation of GCICs as gene-centered regulatory structures rather than predictions of individual transcription factor binding events.

### Gene-level deployment and vocabulary usage of motif families in GCIC islands

To characterize how cis-regulatory islands identified by the GCIC framework are deployed across genes, we next examined motif-family usage patterns at both the gene and island levels (Figure 4). Across all annotated genes, 49.9% contained at least one GCIC island, whereas the remaining genes lacked GCIC-defined regions (Figure 4A). When motif-family occurrences were summed across all GCIC islands associated with each gene, the total motif-family usage was strongly concentrated at low values, with a smaller number of genes exhibiting higher usage, indicating heterogeneity in the extent to which genes are associated with GCIC-mediated regulatory sequence structure.

**Fig. 4.**
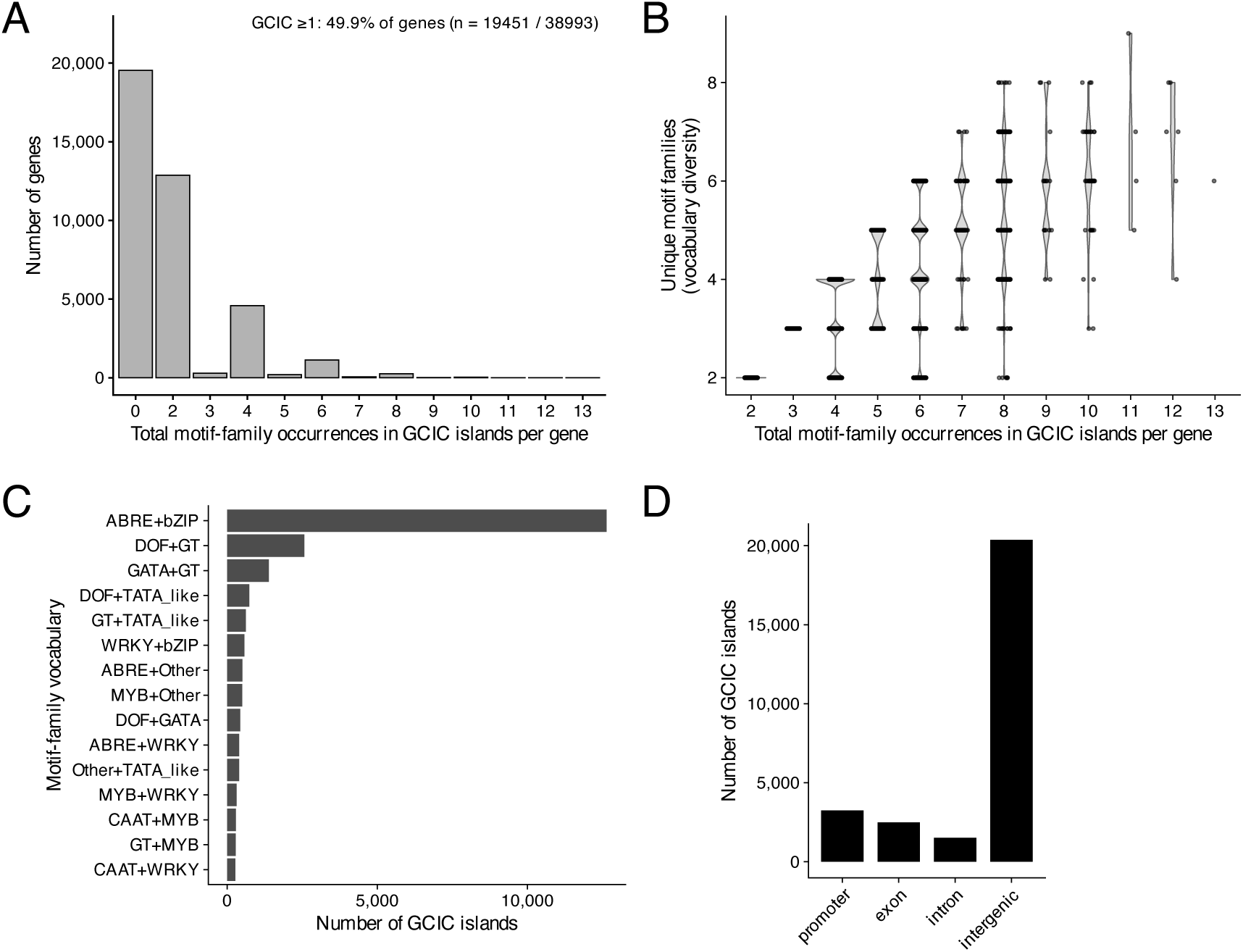
| Gene– and island-level patterns of motif-family usage in GCIC islands. (A) Gene-level distribution of motif-family usage in GCIC islands. All genes were analyzed by summing motif-family occurrences across all GCIC islands associated with each gene. Genes lacking GCIC islands were counted as zero. Overall, 49.9% of genes contained at least one GCIC island. The x-axis indicates the total number of motif-family occurrences per gene, including repeated occurrences of the same motif family across multiple GCIC islands. The y-axis shows the number of genes with the corresponding motif-family occurrence count. (B) Regulatory vocabulary diversity as a function of motif-family usage in GCIC islands. The xaxis, as in Figure 4A, indicates the total number of motif-family occurrences per gene summed across all associated GCIC islands. The y-axis shows the number of unique motif families per gene, which was used as a measure of regulatory vocabulary diversity. Violin plots depict the distribution of diversity values, with individual genes overlaid as jittered points. (C) Motif-family vocabulary usage across GCIC islands. Each GCIC island was classified according to the unique combination of motif families it contains, referred to as a motif-family vocabulary. The bar plot shows the frequency with which each motif-family vocabulary is observed across GCIC islands. For clarity, only the 15 most frequent motif-family vocabularies are shown; the complete list of all vocabularies and their frequencies is provided in Supplementary Table S3. (D) Genomic annotation of GCIC island locations. GCIC islands were classified according to their genomic context, including promoter, exon, intron, and intergenic regions. The bar plot shows the number of GCIC islands overlapping each annotation category, providing a genomic overview of GCIC island distribution.

We next asked how motif-family usage relates to regulatory vocabulary diversity at the gene level. Genes were grouped according to the total number of motif-family occurrences across their associated GCIC islands, and for each gene the number of unique motif families was quantified. As motif-family usage increased, genes tended to harbor a larger number of distinct motif families (Figure 4B). This relationship indicates that increased motif-family usage at the gene level is accompanied by expansion of regulatory vocabulary diversity, rather than repeated use of a fixed subset of motif families.

To examine how motif families are combined within individual GCIC islands, we classified each island according to its motif-family vocabulary, defined as the unique combination of motif families present within that island. Analysis of motif-family vocabulary usage revealed a highly uneven distribution across GCIC islands (Figure 4C). A small number of vocabularies were observed frequently, whereas many others occurred at low frequency (Supplementary Table S3), indicating that GCIC islands collectively exhibit a broad range of motif-family combinations with strongly skewed usage frequencies.

Finally, we examined the genomic context in which GCIC islands are located. GCIC islands were distributed across promoter, exonic, intronic, and intergenic regions (Figure 4D), indicating that GCIC-defined regulatory sequence organization is not restricted to promoter regions or other canonical regulatory compartments. Together, these results characterize GCIC islands as gene-centered regulatory structures that vary widely across genes, with GCIC islands observed in multiple genomic contexts.

### Positioning GCICs within the known cis-regulatory landscape

To contextualize GCICs relative to previously defined cis-regulatory annotations, we quantified their genomic overlap with conserved noncoding sequences (CNSs), dREG peaks, and small RNA‒associated genomic features using publicly available datasets previously described by Goliasse et al. (2025). As a reference frame for interpreting overlap-based analyses, we first examined the extent of overlap among these annotation layers themselves (Figure 5A). Pairwise overlap was limited, and triple overlap among all three layers was rare, comprising only 59 genomic locations across the genome. When calculated relative to each annotation layer, this corresponds to approximately 0.061% of CNSs, 0.182% of dREG peaks, and 0.161% of small RNA–associated genomic features, indicating that multi-layer concordance among existing annotations is uncommon.

**Fig. 5.**
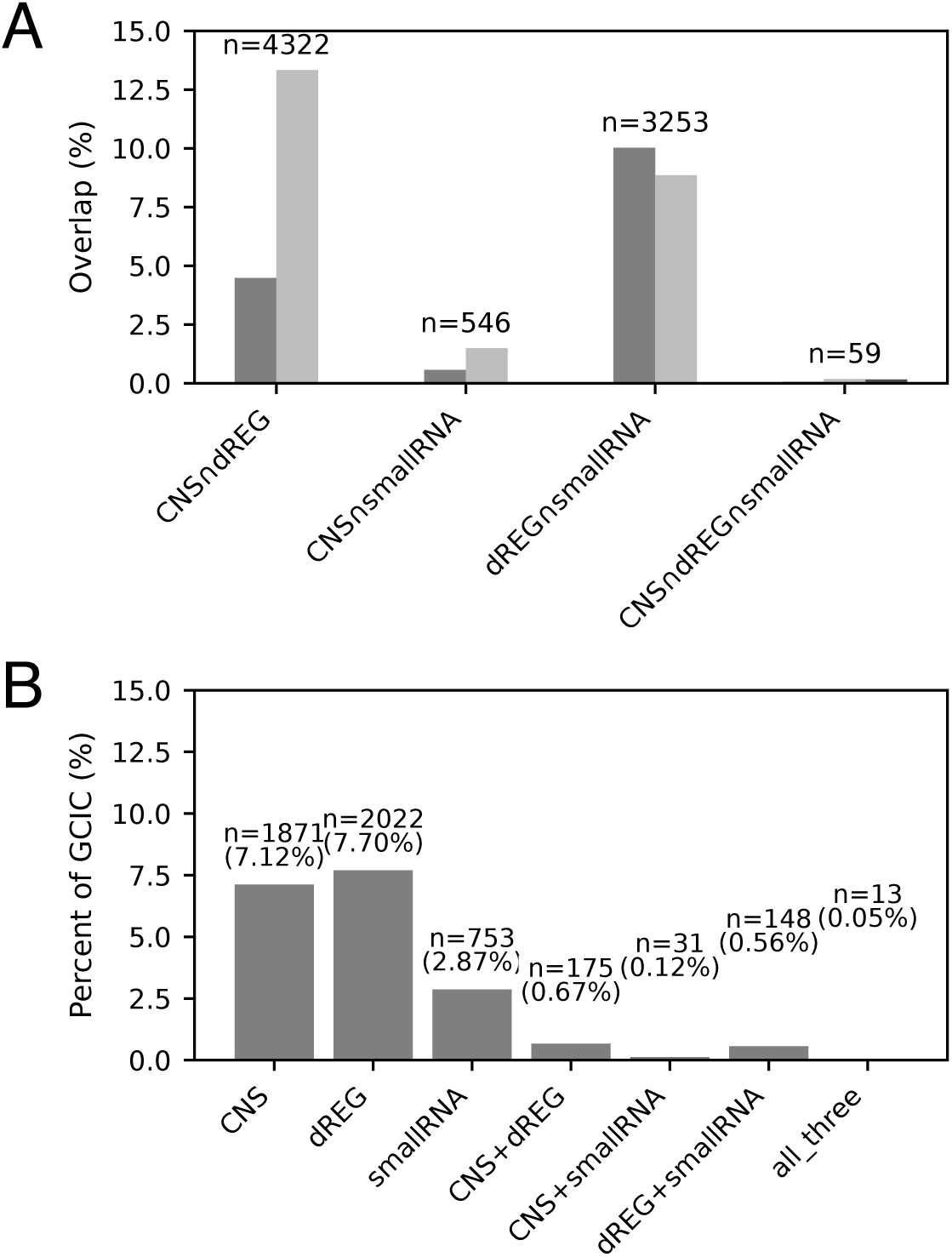
| Positioning GCICs within the known cis-regulatory landscape. (A) Pairwise and three-way genomic overlap among conserved noncoding sequences (CNSs), dREG peaks, and small RNA loci in the Azucena genome. Pairwise overlap among annotation layers is limited, and three-way overlap is rare (59 loci; corresponding to 0.061% of CNSs, 0.182% of dREG peaks, and 0.161% of small RNA loci). (B) Partitioning of GCICs by exclusive and combinatorial regulatory layer support, shown as the percentage of GCICs overlapping CNSs, dREG peaks, small RNA loci, or their combinations. GCICs lacking overlap with any annotation layer are excluded from this panel; their prevalence is described in the main text. Single-layer–supported GCICs constitute the majority of annotated loci, whereas GCICs supported by multiple regulatory layers represent a small, highly selective subset with convergent regulatory evidence.

Consistent with previous qualitative observations that distinct functional features in the rice cis-regulatory landscape often occupy separate genomic locations (Goliasse et al., 2025), our analysis provides a quantitative assessment of pairwise and three-way overlap among these layers.

We then assessed the overlap between GCICs and established cis-regulatory annotation layers by partitioning GCICs according to their layer support (Figure 5B). GCICs overlapped with CNSs and dREG peaks at levels comparable to the overlap observed among existing annotations, whereas overlap with small RNA ‒ associated genomic features was more limited. Partitioning GCICs by layer support revealed that the majority of GCICs were not annotated by any single layer, while progressively smaller subsets were supported by one or multiple layers (Figure 5B).

## Discussion

### Limitations of motif-centric approaches under gene-centered conditions

PWM-based motif scanning and clustering approaches have been highly effective in genome-wide analyses, where background normalization suppresses ubiquitous sequence patterns and enhances detection of TF-specific binding events (Stormo, 2000; Wasserman and Sandelin, 2004; Spitz and Furlong, 2012). However, as illustrated by the gene-centered comparison in Figure 3, removal of genome-wide normalization exposes a strong dependence of these methods on local sequence composition.

In the present gene-centered analytical conditions, FIMO reports motif-like sequence occurrences that are broadly distributed and largely reflect local sequence composition rather than discrete regulatory units. Cluster-Buster, which aggregates dense PWM hits, is similarly sensitive to low-complexity and repetitive sequences when genome-wide background correction is absent. Importantly, this behavior does not indicate methodological failure; rather, it reflects faithful reporting of motif-derived signals under conditions that differ from those for which these tools were originally optimized (Frith et al., 2003; Grant et al., 2011). These observations underscore the extent to which motif-centric approaches rely on genome-wide statistical frameworks to separate regulatory signal from compositional bias.

PLACE-derived regulatory motifs include many short and degenerate sequence elements, typically on the order of 4–8 bp, that are experimentally associated with regulatory activity but do not necessarily correspond to high-information-content transcription factor binding sites (Higo et al., 1999). Under gene-centered constraints, PWM-based methods lack mechanisms to fully exploit the regulatory information encoded in such sequence vocabularies, particularly when regulatory diversity must be distinguished from simple compositional repetition. As a result, motif-centric analyses may conflate regulatory organization with low-complexity sequence accumulation when genome-wide assumptions are relaxed (Tautz et al., 1986; Hardison and Taylor, 2012).

### Cis-regulatory islands identified by a TF-independent, gene-centered framework

The concordance observed between GCIC-defined cis-regulatory islands and experimentally characterized regulatory regions should not be interpreted as validation of GCIC as a predictive tool for transcription factor binding or regulatory activity. Rather, such concordance indicates that GCIC captures aspects of regulatory sequence organization that are already implicit in well-studied loci, but that are not explicitly represented in motif-centric or TF-centric analytical frameworks.

In addition to the intronic regulatory region of *DL*, similar positional correspondence was observed at other rice loci for which cis-regulatory regions have been functionally delineated, including *OsMADS1* and *SiR*. Notably, these loci differ substantially in gene structure, regulatory context, and the experimental paradigms used to define their regulatory regions. Nevertheless, in each case, GCIC-defined islands coincide with genomic intervals that have been shown to be necessary for proper spatial regulation.

Importantly, these observations do not imply that GCIC identifies regulatory elements with base-pair precision, nor that all GCIC islands correspond to functionally validated cis-regulatory modules. Instead, they support the interpretation that GCIC delineates gene-centered regulatory landscapes—extended sequence environments in which regulatory information is organized—within which transcription factor binding and regulatory activity are subsequently realized. From this perspective, transcription factors act upon pre-existing regulatory sequence landscapes, rather than defining regulatory structure de novo, consistent with conceptual models proposed in prior studies (Levine and Tjian, 2003; Spitz and Furlong, 2012).

The observation that GCIC islands can coincide with experimentally characterized regulatory regions across multiple loci, while remaining agnostic to transcription factor identity and genome-wide background normalization, underscores the conceptual distinction between identifying regulatory structure and predicting regulatory activity. GCIC is designed to address the former by abstracting cis-regulatory organization as a property of local sequence vocabulary usage conditioned on individual genes.

### Implications for gene-centered regulatory analysis

Genome-wide analysis of GCIC islands further reveals an apparent tension between regulatory diversity and combinatorial reuse (Figure 4). At the gene level, increased motif-family usage is associated with expansion of regulatory vocabulary diversity, indicating that genes deploy increasingly diverse regulatory sequence vocabularies as GCIC-associated motif usage increases. In contrast, at the level of individual cis-regulatory islands, motif-family vocabularies exhibit strongly skewed usage frequencies, with a limited number of motif-family combinations recurring frequently across islands. Together, these observations indicate that regulatory diversity arises through gene-specific deployment and reuse of a restricted set of motif-family combinations, rather than through unrestricted combinatorial complexity at the level of individual regulatory elements.

Although motif-family vocabularies are derived from the PLACE database, the observed skew in vocabulary usage cannot be trivially attributed to motif frequency bias alone. If database composition were the primary determinant, increased motif-family usage would be expected to result in repeated use of the same motif families rather than the observed expansion of regulatory vocabulary diversity at the gene level. Instead, skew emerges specifically at the level of motif-family combinations, suggesting that higher-order organization of regulatory sequence vocabulary contributes to GCIC island structure.

By conditioning enrichment on individual genes rather than genome-wide averages, identification of cis-regulatory islands using the GCIC framework reveals gene-specific regulatory landscapes that are obscured by global normalization strategies. Genome-wide application of this framework demonstrates that cis-regulatory islands are widespread but heterogeneous, varying in number, genomic position, and motif family composition across genes.

Additional cis-regulatory islands detected outside experimentally validated regulatory regions may reflect latent, redundant, or context-dependent regulatory structures that are not revealed under specific experimental conditions. Together, these findings highlight the importance of aligning analytical assumptions with biological scale and demonstrate that gene-centered identification of cis-regulatory islands provides a complementary, TF-independent lens for interrogating regulatory organization. Similar considerations apply to deep learning–based regulatory models, which optimize predictive accuracy across genome-wide training data but do not explicitly define gene-centered regulatory structure.

### Positioning GCICs relative to established cis-regulatory annotations

The positioning of GCICs relative to established cis-regulatory annotations further clarifies the conceptual scope of the GCIC framework. Using previously published CNS, dREG, and small RNA annotations as reference layers, we show that GCICs are neither trivial rediscoveries of known regulatory elements nor arbitrary sequence features. Instead, GCICs overlap with established cis-regulatory annotations at levels comparable to the overlap observed among these annotations themselves (Figure 5).

Importantly, triple overlap among CNSs, dREG peaks, and small RNA ‒ associated genomic features is exceedingly rare, comprising only 59 genomic locations genome-wide. This provides a stringent reference frame for interpreting multi-layer support. Within this context, GCICs supported by multiple annotation layers represent a highly selective subset, consistent with the interpretation that GCICs capture localized regulatory environments rather than specific regulatory modalities. Conversely, the large fraction of GCICs that do not overlap any single annotation layer suggests the presence of latent or context-dependent regulatory structures that are not readily detected by assays optimized for specific regulatory signals.

These observations reinforce the view that GCICs should be interpreted as indicators of regulatory permissiveness rather than as predictions of regulatory activity under any particular condition. By abstracting cis-regulatory organization in terms of motif-family vocabulary structure, the GCIC framework complements functional genomics approaches that assay specific molecular readouts, providing an orthogonal perspective on regulatory landscape organization.

## Conclusion

In this study, we present the gene-centered identification of cis-regulatory islands (GCIC) as a transcription factor–independent framework for characterizing cis-regulatory organization under gene-centered conditions. By conditioning enrichment on individual genes rather than on genome-wide averages, GCIC identifies localized regulatory regions defined by the coordinated enrichment of diverse regulatory sequence vocabularies within the sequence context of individual genes. Comparative analyses demonstrate that GCIC captures spatially confined regulatory structure that is distinct from both diffuse motif density profiles and broad cis-regulatory modules identified by motif-centric approaches, reflecting differences in underlying modeling assumptions rather than methodological failure.

Positioning GCICs within the broader cis-regulatory landscape further reveals selective but non-trivial correspondence with conserved and functional regulatory annotations, indicating that cis-regulatory islands are neither reducible to existing annotation classes nor attributable to stochastic sequence features. Accordingly, GCICs should not be interpreted as predictions of specific transcription factor binding events, but rather as indicators of regulatory potential—sequence environments that pre-exist and constrain transcription factor action. Together, these findings establish GCIC as a complementary lens for interrogating gene-centered regulatory architecture and provide a conceptual foundation for integrating regulatory sequence organization with transcription factor occupancy, epigenomic state, and evolutionary conservation in future studies.

## Supporting information

Supplementary Data and Tables (GCIC analysis)

## Acknowledgements

This work was supported by the Cabinet Office, Government of Japan, Moonshot Research and Development Program for Agriculture, Forestry and Fisheries (funding agency: Bio-oriented Technology Research Advancement Institution), Grant Number JPJ009237.

